# Online detection of auditory attention with mobile EEG: closing the loop with neurofeedback

**DOI:** 10.1101/218727

**Authors:** Rob Zink, Stijn Proesmans, Alexander Bertrand, Sabine Van Huffel, Maarten De Vos

## Abstract

Auditory attention detection (AAD) is promising for use in auditory-assistive devices to detect to which sound the user is attending. Being able to train subjects in achieving high AAD performance would greatly increase its application potential. In order to do so an acceptable temporal resolution and online implementation are essential prerequisites. Consequently, users of an online AAD can be presented with feedback about their performance. Here we describe two studies that investigate the effects of online AAD with feedback. In the first study, we implemented a fully automated closed-loop system that allows for user-friendly recording environments. Subjects were presented online with visual feedback on their ongoing AAD performance. Following these results we implemented a longitudinal case study in which two subjects were presented with AAD sessions during four weeks. The results prove the feasibility of a fully working online (neuro)feedback system for AAD decoding. The detected changes in AAD for the feedback subject during and after training suggest that changes in AAD may be achieved via training. This is early evidence of such training effects and needs to be confirmed in future studies to evaluate training of AAD in more detail. Finally, the large number of sessions allowed to examine the correlation between the stimuli (i.e. acoustic stories) and AAD performance which was found to be significant. Future studies are suggested to evaluate their acoustic stimuli with care to prevent spurious associations.

## 1 Introduction

Understanding speech within a noisy or multiple-speaker environment is one of the auditory system’s most difficult tasks. Nevertheless, humans have the astounding capacity to focus their attention to interesting sources of speech within complicated audio scenes. This allows listening to one speaker of interest while simultaneously ignoring other speakers and sound sources competing for attention. This problem was coined the ‘cocktail party problem’ by Colin Cherry in 1953 [10]. The remarkable capability of the human brain to solve this problem seemingly easy motivated researchers to unravel the neural underpinnings to this phenomenon. Significant progress came from the finding that cortical oscillations phase lock to the envelope of the speech stream [1]. Consecutively, attended speech was found to be more prominently represented in electrophysiological signals as compared to unattended speech, especially at higher cognitive levels [23, 13, 20]. [32] illustrated that lower auditory areas of the brain encode both attended speech and background noise, whereas higher auditory areas have a preference for the attended speech stream only. Based on these findings it became possible to detect which speaker (i.e. in a two-speaker cocktail party scenario) was attended to from the recorded activity from electroencephalography (EEG), magnetoencephalography (MEG) and cortical surface recordings [29, 2, 23]. This process will be referred to as auditory attention detection (AAD). These models followed mostly two approaches: define a regression model that predicts the neural responses based on the sound, so-called forward modeling [4, 2, 3], or reconstruct the speech envelopes of the (un)attended speaker from the EEG or MEG neural data (backward modeling) [29, 11]. Others have experimented with alternative approaches, for example to extract features from the cross-correlation of neural response and speech envelope to train a linear classifier [19].

In this paper we focus on EEG, which has been employed successfully for such auditory attention detection (AAD). More specifically on the backward modelling of AAD; a pretrained decoder that makes a linear combination of the EEG and delayed versions of it allows to extract a signal with a significantly higher correlation to the attended speaker’s speech envelope as compared to the unattended speaker’s envelope [29]. In such dual-speaker scenarios, the subject’s attention to a specific speaker could be reliably detected with high accuracy in research labs (e.g. [29]). In the past years several aspects of the analysis pipeline have been optimized to enhance the estimation made by the decoder to increase detection accuracy. Different ways of decoder construction and envelope extraction methods have been evaluated [9, 14] whereas others focused on removing lateralization bias in the decoder and improving stimulus preprocessing to enhance the realism of the experiments [11]. Whereas most AAD studies assume availability of the clean speech envelopes, also more realistic scenarios have been studied where these per-speaker envelopes are not available. Successful extraction of per-speaker envelopes from noisy speech mixtures by means of blind source separation techniques has been demonstrated in [35], where AAD was used to steer an acoustic beamformer towards the attented speaker. Alternatively, deep neural networks have been employed to obtain a reliable AAD system based on mixed audio input [28]. Furthermore, the influence of noise on the audio waveforms used in the decoder was rendered insignificant to a certain extent [6]. Another study provided evidence that only a specific set of spatial regions is involved in the detection process and consequently the number of electrodes may be decreased drastically (e.g. from 96 down to 15) and still achieve robust classification accuracy, which would be especially meaningful for mobile experiments in which there are only limited number of electrodes available [26]. [17] showed that the focus of attention on one of two simultaneously presented speech streams was not influenced by including additional competing speakers from different spatial locations.

These studies on auditory attention detection were conducted in a lab setting, using clinical grade equipment. However, when considering the development of consumer-geared applications, the system needs to be robust to daily-life settings. A recent study addressed this issue by providing proof-of-concept AAD using EEG electrodes positioned around the ear (i.e. cEEGrids) that would allow for mobile AAD, although the classification accuracy was lower as compared to a traditional EEG cap, potentially due to suboptimal electrode placement [25]. Another recent study reported that the EEG response from a single in-ear electrode, hearingaid-compatible configuration provides valuable information to identify a listeners focus of attention [16]. However, application of a true online EEG-based AAD system has not yet been evaluated to date.

The effectivity of EEG-based AAD depends not only on the temporal resolution and the acoustic scenario, but also on the responses of the test subjects themselves (e.g. real and pseudo BCI illiteracy, see [38]). Large differences in accuracy between subjects were observed in the aforementioned studies. It is known for other cognitive paradigms (e.g. such as P300 oddball studies [41]) that subjects’ physiological responses can differ substantially, even within subjects when moving from restricted to more real-life scenarios [41]. Providing feedback about the ongoing EEG signals was shown to be beneficial for other EEG paradigms to strengthen the brain responses (e.g. P300 ERP [7]) or induce specific neural patterns (e.g., motor imagery [37]). A similar reasoning could be applied for AAD, i.e. training users to elicit relevant brain responses with respect to the attended speech stream might increase the accuracy. To this end users might experience positive effects from such a feedback training to increase the performance of EEG-based AAD.

Changing the brain’s neuronal activity in order to gain rewards is basically operant conditioning of neural activities. This is often referred to as neural biofeedback, or neurofeedback (NFB) whenever the brain activity of a human is the target. Neurofeedback training (e.g. through fMRI or EEG) has been defined as a method to self-regulate one’s own brain activity [15, 33]. Adding a feedback in the typical BCI or AAD analysis closes the loop; measurements of neural activity are taken, and a real time representation of it is shown whether tactile, auditory or visual to the subject, thus facilitating neural activity self-regulation. Users can learn to control various aspects of their neural activity through a training process which displays online changes of relevant EEG features [27]. NFB training thereby engages several learning mechanisms in the brain of which operant conditioning is considered as the main principle [27, 15]. Changes in brain activity that reflect successful NFB training should be rewarded or positively reinforced. If the represented performance measure reaches a certain threshold, a rewarding stimulus should be presented. Typical visual feedback stimuli are a moving cursor, a bar or sphere varying in size or objects changing in color. Feedback is often presented in a game-like format. [27]. Such formats intend to increase and maintain the subject’s motivation and attention, which helps facilitating self-regulation of neural activity and at the same time could cause increased behavioral task performance (i.e. create a positive state of mind). However, besides the behavioral control-input task, the subject thus has a second task to reach a specified goal regarding the NFB presentation. This results in a higher cognitive load and a division of cognitive resources between both tasks, which could lead to poorer BCI performance [21]. Therefore, the feedback signal should not distract the subject too much from the behavioral task to be performed [27]. For an in-depth review of the theories and models that have been proposed to explain the neural mechanisms of neurofeedback learning, we kindly refer the reader to a recent review by Sitaram et al. [33]. Training with neurofeedback has been utilized as a therapy tool for the normalization of brain activity, or as a tool for cognitive enhancement in the case of healthy subjects [15]. For example, NFB training for the improvement of attention ability [31], improvement of ADHD (attention deficit hyperactivity disorder) treatment [5], epilepsy [34] and post-stroke motor learning (i.e. through motor imagery (MI)) [37]. The latter furnished a proof-of-concept for a mobile, low-density EEG system for effective NFB training at the home of the participant. Furthermore, recent case study reports by [39] strengthened the notion that MI training at home can lead to noticeable changes in brain patterns. Considering the AAD paradigm there is no reference study or standardized training protocol to date. Hence the method and extent that self-regulation of the neural responses can be applied to such an auditory attention task (i.e. speaker detection) remains unknown. Nevertheless, it should be noted that few reports exist of self-regulation of activity from human auditory areas facilitated by NFB training (e.g. slow cortical potentials [24] and fMRI based [36]).

The aim of the current work is twofold. Firstly we demonstrate the application of the AAD in a real-time setup. Twelve subjects were recorded in an office environment with mobile EEG hardware and stimuli were presented through consumer-grade headphones. In addition, we apply a neurofeedback scenario to provide a proof-of-concept for a fully automated closed loop AAD system. We implemented an online AAD system with a time resolution of 10 seconds with a backward decoding approach. Secondly, longitudinal data was collected from two subjects (of which one received the neurofeedback) to evaluate the feasibility of longterm neurofeedback training on AAD. The EEG was recorded at home in a 24 minute session for seven days (spread over four weeks). Note that although we report a case study a large number of sessions were recorded over time allowing to investigate longitudinal effects that have not been investigated in the literature to date. Together, these studies describe the feasibility of in home AAD recording and the potential of online AAD feedback for improving the AAD abilities of humans. In this setting we decoded the subjects’ attention with high accuracy and report on the influential factors for training of the AAD protocol. In addition, we highlight aspects in the AAD experimentation that need deliberation in order to validate and compare AAD results between studies. These results pave the way to further investigate how subjects might be able to increase the AAD performance and thus improve the applicability of AAD in general.

The paper describes both studies in parallel. Unless specified separately, both studies share the same specifications. The initial experiment that was used to setup the closed-loop online system will be denoted as ’Study I’ and the in-home longitudinal case study as ’Study II’. In general the paper is structured as follows: the most important experimental details and setup are described in the first part of the methods section, followed by a description of the online feedback procedure (which deviated slightly between the two studies). The results section firstly reports on the initial study (study I) with emphasis on the closed-loop system and neurofeedback presentation. The latter part of the results section highlights the findings related to longitudinal AAD training effects (over multiple days) and the role of the audio stimuli used in the current study (study II). Finally, results of both studies are considered in perspective and future perspectives are outlined.

## 2 Methods

### 2.1 Participants

Twelve native Dutch-speaking subjects (mean age (SD) 22.4 (± 2.1) years, six women) participated in study I. In study II two native Dutch-speaking subjects took part, aged 21 and 26, both female. All subjects reported normal hearing and no past or present neurological or psychiatric conditions. All participants signed informed consent forms prior to participation. The ethics committee of the KU Leuven approved the experimental setups.

### 2.2 Data Acquisition

The acquisitions (study I & II) were conducted with a SMARTING mobile EEG amplifier from mBrainTrain (Belgrade, Serbia, www.mbraintrain.com). This amplifier comprises a wireless EEG system running on a notebook computer (Dell XPS 9550) using a small 24-channel amplifier with similar characteristics to a stationary laboratory amplifier (*<*1uV peak to peak noise; 500Hz sampling rate). The EEG was measured using 24 Ag/AgCl passive scalp electrodes (Easycap), placed according to the 10-20 standard system with positions: FP1, FP2, Fz, F7, F8, FC1, FC2, Cz, C3, C4, T7, T8, CPz, CP1, CP2, CP5, CP6, TP9, TP10, Pz, P3, P4, O1 and O2. Impedances were kept below 10 kOhm and an abrasive electrolyte gel was applied to each electrode. EEG data were recorded through Openvibe [30] (and stored for offline (reference) analysis) and streamed from Openvibe to Matlab via the labstreaminglayer interface (LSL) [22]. The audio stories were played via Openvibe and pre-loaded into Matlab for the online analysis part and synced via the Openvibe audio triggers. Every ten seconds, the data was retrieved from the LSL stream and analyzed in Matlab. Both online and offline analysis used custom-made Matlab scripts.

### 2.3 Stimuli

The audio stimuli of Study I consisted of four fairytale stories in Dutch (of approximately 12 minutes’ length), narrated by four different male Dutch (Flemish) speakers, each story was split in two parts (blocks) of ±6 minutes (8 blocks in total). Study II consisted of 32 blocks (of similar ±6 minute length) of Dutch fairytale stories (not overlapping with Study I) narrated by different Dutch male speakers. A total of 48 min stimuli were prepared for Study I and 192 min for Study II.

Silences in the audio were truncated to 500ms and audio waveforms’ RMS were normalized per story (i.e. 6 minutes) similarly as reported previously by [11]. Subjects listened to two stories presented simultaneously through low-cost consumer headphones (Sennheiser mx475). The two audio streams were filtered by head-related transfer functions leading to more realistic perception [11]. Subjects were asked to pay attention to only one story on the left or right side. After each block, multiple-choice questions were presented and subjects indicated the difficulty in listening (i.e., indicated on a ten-point scale) and likewise their interest in the current story. Post-hoc we evaluated potential relationships between these user metrics and the AAD performance.

### 2.4 Procedure

An overview of the procedures of both studies are schematically illustrated in Figure 1.

**Figure 1:**
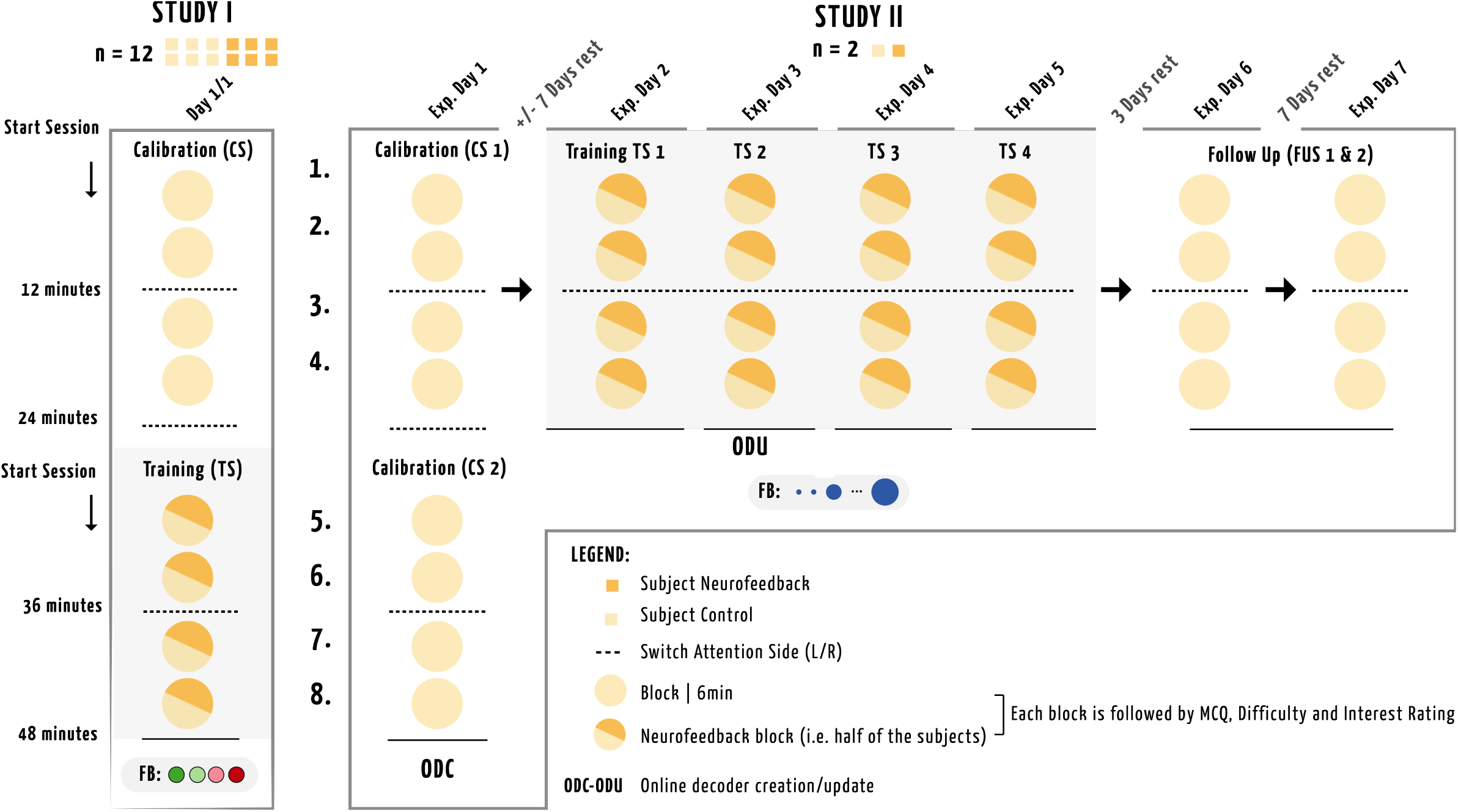
Overview of the recording sessions and days for both studies. Values depicted are for illustrative purposes only. Study I and Study II had different audio stimuli and each circle in the figure indicates one block of 6 minutes. After listening to one full story (2 blocks of 6 minutes), subjects switched attention to the opposite side, i.e. left or right speaker, to avoid introducing a lateralization bias in the decoder. Blocks in the shaded area represent that half of the subjects were presented with visual feedback (different per study) and at these blocks a pre-trained decoder was used based on the data of previous sessions. Study I consisted of two sessions of two stories (four blocks) each. Study II consisted of 8 sessions: 2 calibration (CS), 4 with feedback training (TS) and 2 follow up sessions (FUS).

#### 2.4.1 Study I

The experimental setup consisted of two sessions of two stories (four blocks) each. After listening to one full story (2 blocks of 6 minutes), subjects switched attention to the opposite side (i.e. left or right speaker) to avoid introducing a lateralization bias in the decoder [11]. The first session was used for calculating the decoder (calibration), the latter for evaluation (testing). Half of the subjects received visual feedback (feedback group) on the laptop screen for the second session of the experiment (as described in section 2.6), whereas the other subjects received no feedback (control group). Order effects were avoided by alternating the listening sides among subjects. The study took place in an office–like setting while seated on a chair at a desk.

#### 2.4.2 Study II

The experiment consisted of 8 sessions: 2 calibration (CS), 4 training (TS) and 2 follow up sessions (FUS). Similarly to Study I each session consisted of four 6 min blocks and ear attendance was switched from right to left at mid-session. The calibration sessions were used to calculate the decoder that could be used for online detection during training session 1&2. A subject-specific decoder was constructed for both subjects (to allow for equal comparison of the subjects’ results). This online decoder was updated one time, for both subjects, after training session 2, based on the subject-specific data collected during training session 1&2. During training phase, one participant, further referred to as SF (i.e. Subject Feedback), was presented with feedback about ongoing AAD performance (see section 2.6). The other participant, SC (i.e. Subject Control), did not receive any feedback throughout the entire study, but participated in as many sessions as SF. During follow up sessions, neither subject was presented with NFB and subjects’ offline AAD performance was evaluated to assess for long-term training effects. Sessions took place at the subject’s home, in a comfortable seated position in the living room.

### 2.5 Analysis

The following paragraphs provide further details on the closed-loop implementation of the AAD. It describes the preprocessing steps, decoder construction, detection (i.e classification) process and feedback procedure. These stages are schematically depicted in Figure 2.

**Figure 2:**
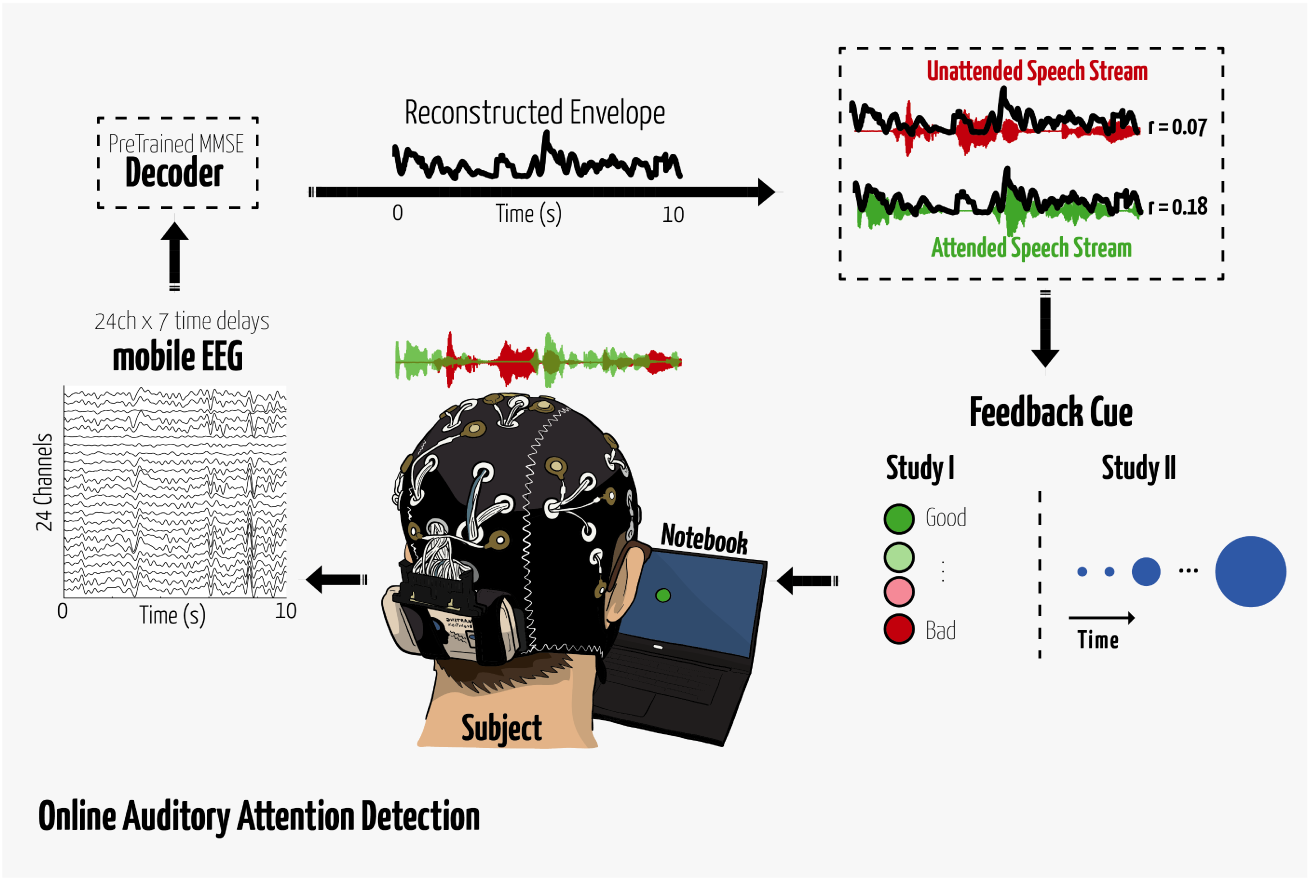
Illustration of the Auditory attention closed-loop analysis. Values depicted are for illustrative purposes only.

#### 2.5.1 Preprocessing

EEG data were bandpass-filtered at 1-8 Hz and consequently down-sampled to 20Hz as implemented in the EEGLAB toolbox [12] for each 10s segment of data. The absolute value of the audio waveforms with power-law compression with exponential 0.6 was taken to obtain the audio envelopes, and then an 8Hz low-pass filter was applied [9]. Envelopes were extracted from the clean audio signals for the separate speakers.

#### 2.5.2 Decoder construction

Detection of auditory attention was based on the EEG-based method of stimulus reconstruction which was first presented by [29]. The goal is to reconstruct the envelope of the attended speech stream *S* based on neural recordings *R* via a linear spatiotemporal decoder *D*. In case of *N* electrodes, *R*(*t, n*) represents the EEG signal recorded at electrode *n* at sample time *t*. A decoder *D*(*τ, n*) maps from *R*(*t, n*) to the attended speech stream envelope *S*(*t*) as follows:

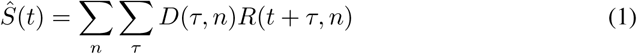

where 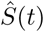 denotes the estimated envelope of the attended speech stream and *τ* represents a time lag. 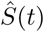 is thus estimated from the EEG recordings at all *N* electrodes and time-lagged versions of those recordings. This way, information in the EEG at later time points is included in estimating the speech stream envelope at earlier time points. The decoder D is computed that minimizes the mean-squared error (MMSE) between the actual and reconstructed envelope. Analytically, *D* is found by:

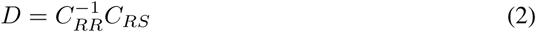

where *C*_*RR*_ is the auto-correlation matrix of the EEG data (across all electrodes and time lags), and *C*_*RS*_ is the corresponding cross-correlation vector between the attended speech envelope and the EEG data (across all electrodes and time lags).

In both studies, six equidistant time-shifted versions of the EEG trial were obtained in a 50-300ms range after stimulus onset, i.e., *τ* ranges from 0 to 6 in equation 1. All EEG channels and their 6 delayed versions are then linearly combined using a pre-trained linear decoder following Equation 2; all the *C*_*RR*_s and *C*_*RS*_ s computed over the training trials were averaged to create a single average covariance matrix and cross-correlation vector [9].

The decoders were trained per subject following an (offline) leave-one-trial-out structure. The decoders for the feedback parts of the experiment were computed solely on the calibration data of the previous session (Study I) or previous days (Study II, see 2.4.2) to allow for online classification. To make topographic plots of the decoder, single per-channel weights were obtained (for each session and for each subject separately) by averaging the decoder coefficients across all time lags which is similar to the procedure reported in [26]. The resulting topoplots are only used to compare decoders across different sessions or subjects, and should not be interpreted individually, as they do not necessarily highlight the activated brain regions, nor the importance of each channel in reconstructing the attended speech envelope. The latter would require an additional correction to take correlations across channels into account (see equations 25-26 in [8]), and the former requires a forward model instead of the backwards decoder used here.

#### 2.5.3 Attention detection

For each test trial Pearson’s correlation coefficient is employed to quantify the decoders’ reconstructed envelope to the attended stimulus (CA) and unattended stimulus (CUA). The highest correlation value determines to which of the two speakers the subject was listening at the current trial. Offline analysis was done on trial lengths from 10s down to 1s. Real-time analysis was done solely on trial lengths of 10s. This trial length was based on results from a small pilot study (i.e. four subjects, not included in the current manuscript) conducted prior to Study I. The pilot study resulted in accuracies *>*80% for trial lengths of 10s which was deemed sufficient to provide accurate feedback to the subjects at an acceptable rate.

### 2.6 Neurofeedback Cue

The predictions of the AAD system were presented online to the subjects such that the feedback cue reflected the degree of which the neural signals of the current trial matched those in the decoder to detect the attended speaker. In our studies, we implemented two different versions of visual feedback on the center of the screen (to minimize eye movement related changes).

#### 2.6.1 Study I

The six subjects who received feedback were presented with a colored circle in the center of the screen. After every test trial of ten seconds, the colored circle indicated the performance of the past ten seconds. Four different colors were used for performance indication: Dark red, light red, light green and dark green from worst to best respectively. Thresholds determining the colors were based on the training set (i.e. first session) in such a way that the CA and CUA difference would be equally divided for the correct and incorrect trials. This means that the median of all positive CA–CUA differences is used as threshold for dark green. Likewise, the median of all negative differences corresponds to the threshold below which the cue is dark red. Note that trials with a larger CA as compared to CUA (i.e., correct trials) are always green (light or dark) and the incorrect trials are always red (light or dark). Decoders for analyzing the neurofeedback sessions (i.e. second half of the data) were based on the individual calibration data as stated in the previous paragraph. Figure 2 illustrates the feedback cues for Study I, only one circle was shown for every trial in the center of the screen.

#### 2.6.2 Study II

After evaluation of the feedback effects of study I, as presented in the Results section 3.1.2, the feedback appearance was modified to avoid potential effects of negative reinforcement penalty; after seeing a light red cue subjects more frequently performed incorrect in the subsequent trial. Therefore, incorrect trials (with CA*<* CUA) were not explicitly visualized in the feedback cue of Study II. Feedback was presented as a blue sphere positioned at the center of the computer screen. In case of above-threshold performance (CA-CUA*>*0), sphere radius grew proportionally to the performance measure (CA-CUA). In case of below-threshold performance (CA-CUA*<*0), the sphere did not grow nor shrink (with the subject not being explicitly informed what this meant). The experimental subject (SF) was instructed to maximize sphere size by focusing on the to-be-attended speech stream. Figure 2 depicts an example of the feedback cues used in this study.

## 3 Results

### 3.1 Study I: Closed-Loop online AAD

#### 3.1.1 Classification Performance

Grand average classification accuracy for the 10s trial length was 81.9% (SD =5.9%). Increasing the window length in the offline analyses to 60 seconds raised the accuracy up to 96.9% (SD=3.5%). All subjects scored above chance level (*>*57%) for the 10s window length.To investigate the potential for providing feedback at a higher temporal resolution we evaluated post-hoc the performance on trial lengths below 10 seconds. Figure 3 shows the classification accuracies for these trial lengths down to 1 second. It can be noted that there is a stronger decrease when moving from 3 to 2 and 1 second window length, although almost 70% of accuracy can be reached on average at a temporal resolution of 3s. For every trial length all subjects scored above chance level; chance levels were corrected for multiple comparisons (i.e. trial lengths) by a Bonferroni correction.

**Figure 3:**
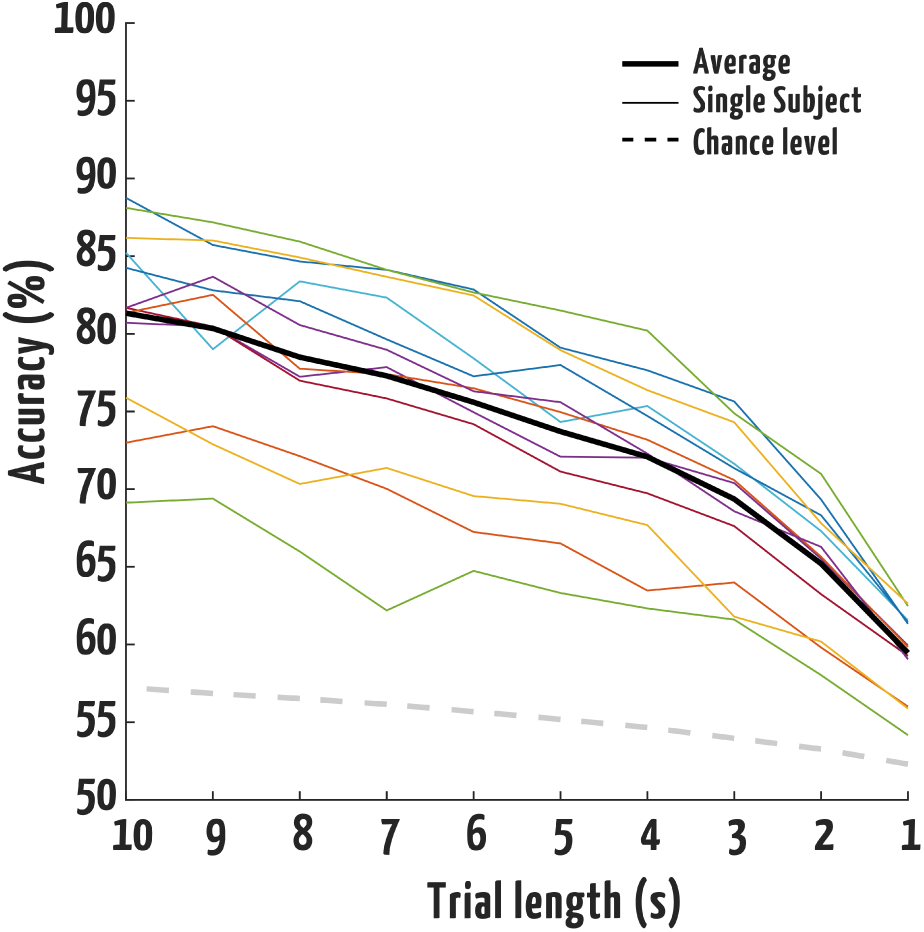
Average and single subject accuracies per trial-length as calculated post-hoc. The dotted line indicates chance level as corrected for multiple comparisons

#### 3.1.2 Neurofeedback Sessions

Average classification accuracy in the online decoding for the second half of the data, (session 2, see Figure 1) was 79.7% (SD = 7.0%) using the pre-trained decoder of session 1. When evaluating the offline leave-one-out decoder for this part of the data, average accuracy was 83.0% (SD = 7.5%). This increase in accuracy was significant (two-sided paired sampled t-test, t11=4.1 p*<*0.01), indicating that training data of the same session resulted in a small classification increase at the 10s windows. Differences in accuracy between the control and feedback group were not evaluated due to the high variability in AAD performance between the subjects (see Figure 3) and small sample size.

Comparing the accuracy within the feedback group between the feedback session and the calibration session revealed a slightly higher accuracy in the feedback session, +4,1 percentage points as calculated with the leave-one-out decoder when averaged over all subjects in the feedback group. Five out of six subjects scored higher in the feedback session, compared to the training session. For the group not receiving feedback, the difference between the second session and the training session was -0.2 percentage points. Surprisingly, no effect was found for increased or decreased CA or CUA changes in the feedback group with respect to the training session.

Relative occurrence of the four feedback cues was on average 41.8% dark green, 40.7% light green, 10.1% light red and 8.7% dark red. We evaluated the average (over subjects) temporal patterns at which the cues were evident in the feedback session. Entries in Figure 4 indicate, by the colors the relative frequency at which a colorcue in the rows was followed by the colorcues in the columns. It can be noted that, in the feedback session, participants shifted more frequently to dark red after seeing light red. Contrastingly, after seeing a dark red cue, the following trial was less likely to be light or dark red - this may indicate awareness of the (very) bad performance and increased effort to compensate, although this remains speculation only. Note that the absolute CA-CUA difference for the light colors is smaller, compared to the dark colors. For example, light green denotes a correct decoding, but with less confidence than in the case of dark green (as described in section 2.6).

**Figure 4:**
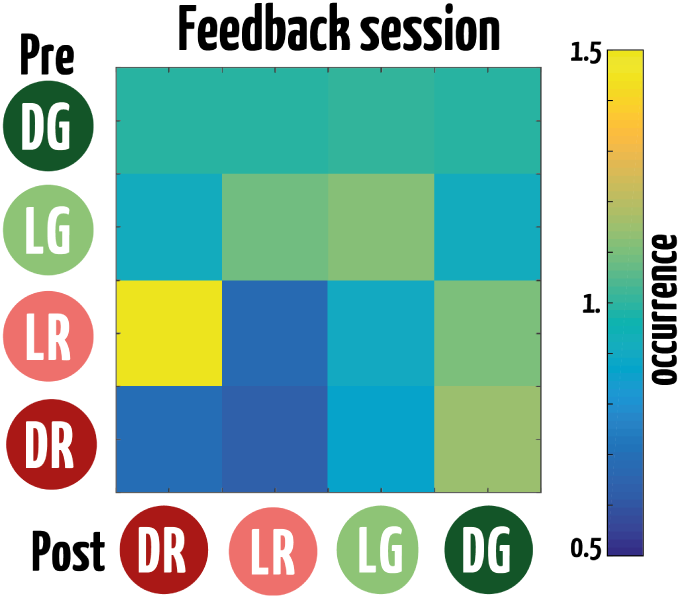
Adjacency matrices for the Neurofeedback session. The entry color indicates the normalized frequencies of a specific color cue (in the rows) that is followed by any other cue (columns). Cues: DG = Dark Green, LG = Light Green, LR = Light Red and DR = Dark Red.

#### 3.1.3 User Metrics

On average, subjects answered 84.2% (SD = 7.9%) of the questions correctly. We contrasted the number of correct responses of each subject to the general accuracy at the 10s window analysis. This revealed a strong positive correlation (r = 0.72, p *<*0.05). One subject was removed, as its number of correct responses differed more than 2 standard deviations from the mean. Figure 5A displays the individual subjects’ accuracy and number of correct responses. A regression line has been added for illustrational purposes. A moderate negative correlation (r = -0.56, p = 0.059) was found between the average accuracy and the subjects’ reported difficulty in answering the questions and overall listening. This correlation is depicted in Figure 5B.

**Figure 5:**
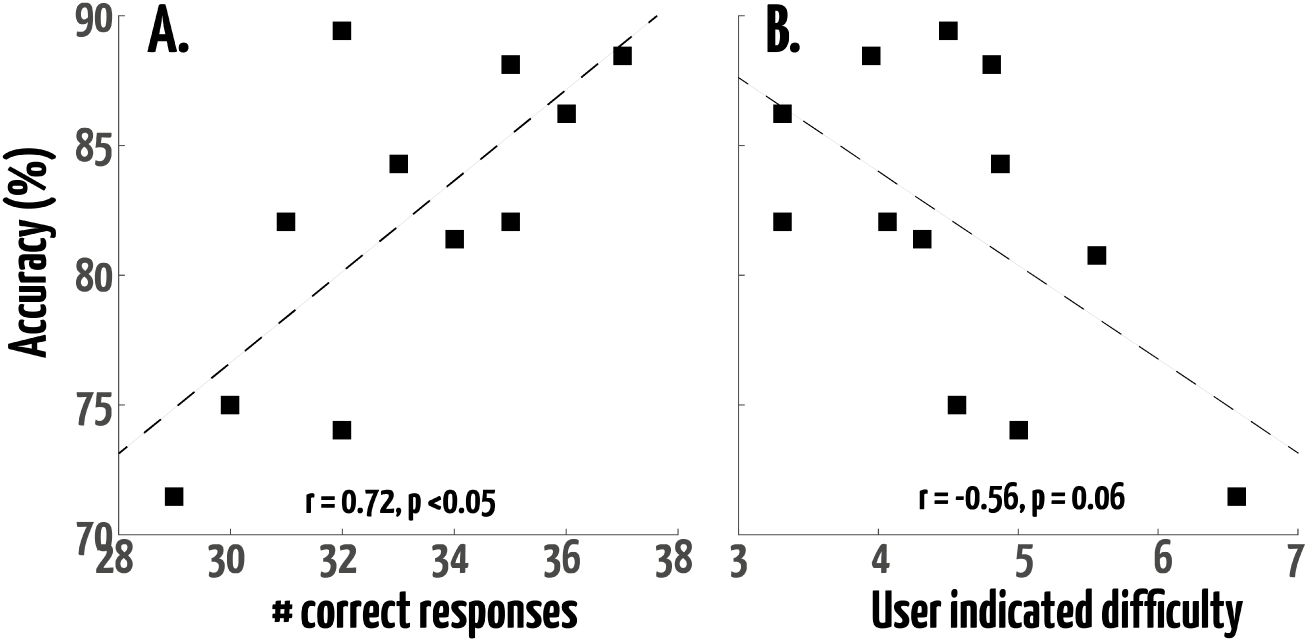
Scatterplots illustrating the correlation between the decoder grand-average accuracy and the number of correct responses after each story in A and the User indicated task difficulty in B. A regression line has been added for illustrational purposes.

### 3.2 Study II: Longitudinal Recordings at home

#### 3.2.1 Classification Performance

Average offline AAD accuracy (and SD) across all days and sessions was 79.7 (± 3.9%) for SC and 79.0 (± 5.9%) for SF. Average online AAD accuracy across training sessions was 78.3 (± 2.6%) for SC and 79.2 (± 2.2%) for SF. Table 1 shows average and range of online AAD accuracies during the training at a block and session level, for both subjects.

**Table 1:** Online AAD accuracies on training blocks and training sessions: range and AVG (± SD) for both subjects.

| ONLINE | | MIN | MAX | AVG±SD |
| --- | --- | --- | --- | --- |
| **Block** | SC | 72.2 | 88.9 | 78.3±5.7 |
| | SF | 66.7 | 88.9 | 79.2±7.0 |
| **Session** | SC | 75.7 | 80.6 | 78.3±2.6 |
| | SF | 77.1 | 82.0 | 79.2±2.2 |

Figure 6A shows online accuracy during training sessions (TS) and offline accuracy during calibration (CS) and follow up (FUS) sessions, for both subjects. Dashed lines indicate the on- and offset of the presentation of NFB to SF (left and right, respectively). No consistent performance increase can be noted for either subject during the training days. These results would indicate the absence of a training effect (regardless of the neurofeedback). However, a strong correlation was found between the session performances of both subjects (r=0.81, P*<*0.05). This illustrates that a potential training effect might be masked by differences in the audio stimuli across sessions. Paragraph 3.2.4 highlights these results more in detail.

**Figure 6:**
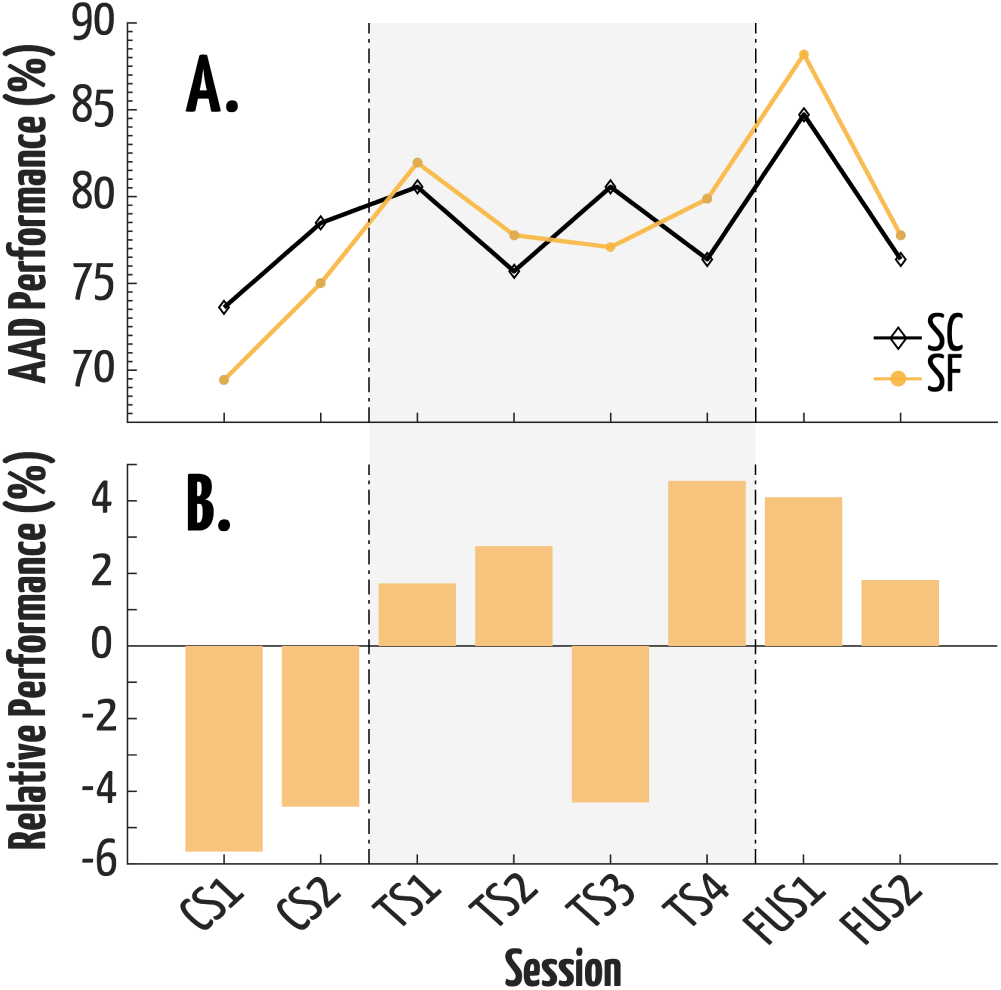
Evolution of a) session performance for both subjects and b) relative session performance of SF to SC. Dash lines indicate the on- and off-set of NFB (shaded area) to SF (left and right, respectively).

Comparing the relative performance of SF to SC removes the potential influence of the audio stimulus, and revealed a more specific effect of the feedback (Fig. 6B). While SF performed almost 6% worse than SC initially, SF became increasingly better than SC throughout training sessions (except for TS3). Furthermore, relative performance remained increased after training (at FUS1), but diminished over time.

Relatively to SC, SF did not show a consistent increase in mean CA-CUA throughout training sessions. Figure 7 shows CA-CUA variance per session, for both subjects. Both subjects started with similar level of CA-CUA variance. While variance of SC was fluctuating around the initial variance, the variance of SF was consistently lower from the training sessions onwards. Relative CA-CUA variance between the two subjects showed the exact inverse pattern of relative performance: consistently decreasing throughout training sessions (from -38% TS1 to -50% at TS4, with inconsistency at TS3), with a partly remained effect at FUS1 (-40%) that diminished towards FUS2 (-35%) (c.q. figure 7 and 6B).

**Figure 7:**
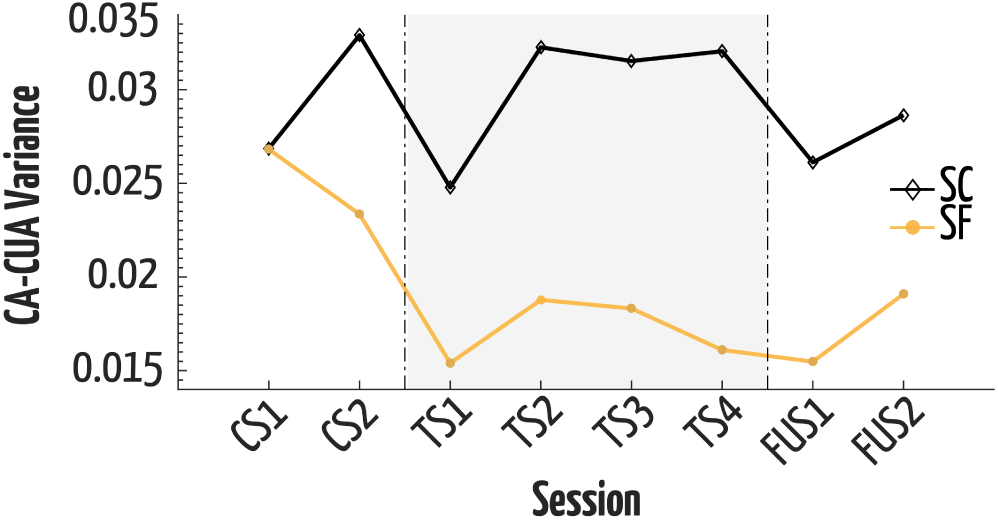
Evolution of CA-CUA variance for both subjects. Dashed lines indicate the on- and off-set of NFB to SF (left and right, respectively).

### 3.2.2 Decoder Characteristics

Figure 8 depicts the spatial distribution of decoder weights (averaged across time lags, see 2.5.2) during the experiment, for both subjects separately. These topoplots show the decoder pattern as calculated on the indicated sessions. For visual comparison throughout the study, we averaged topographies per 2 sessions, this way the effect of the online decoders can be visualized. Note that during TS1&2 feedback was shown with a decoder based on CS1&2 and likewise TS3&4 with a decoder based on TS1&2 (See paragraph 2.4.2). Blue and red color indicate relatively low and high electrode weight, respectively. Note that colors are only relative within a topographic map, and dashed lines indicate the on- and offset of the presentation of NFB (shaded area) to SF (left and right, respectively). The spatial distribution of SF is more consistent as compared to SC throughout the experiment as is evident through higher and more stable correlations. For SF correlations between CS1&2- TS1&2 (r=0.66, P*<*0.001), TS1&2-TS3&4 (r=0.76, p*<*0.001) and TS1&TS2-FU1&FU2 (r=0.72, p*<*0.001) were deemed highly significant. These values indicate that during TS3&4 and to a lesser extent TS1&2 the subjects’ spatial distribution resembled closely the decoder’s pattern. During the follow-up sessions (FU1&2) patterns were still highly similar to the training days. These distributions for SC seem less consistent than that of SF based on lower correlations throughout the experiment resulting in lower correlations which were to a lesser extent (marginally) significant, CS1&2-TS1&2 (r=0.30, p=0.15), TS1&2-TS3&4 (r=0.40, p=0.51) and TS1&TS2-FU1&FU2 (r=0.57, p*<*0.05)).

**Figure 8:**
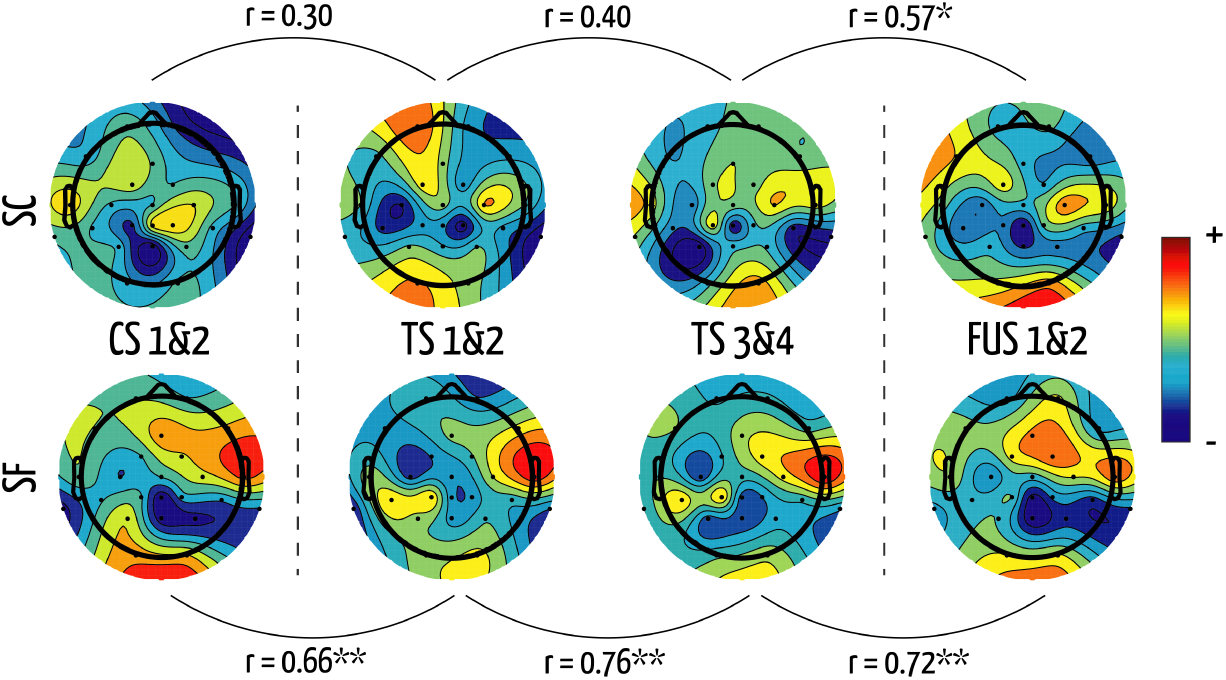
Topographic maps of the spatial distribution of decoder weights throughout the experiment, for a) SC and b) SF. (Red: high weight. Blue: low weight). Dashed lines indicate the on- and offset of NFB to SF (left and right, respectively). Correlations between consecutive spatial distributions are indicated, for both subjects. * denotes p<0.05 and ** p<0.001

#### 3.2.3 User Metrics

On average, SC correctly answered 86.7 (SD = ± 6.2%) of the multiple-choice questions that were asked during a session. For SF, this was 89.8 (± 5.7%). These values indicate that the subjects were successful in following the attended story. Average interest ratings per session were 5.8 (± 0.8) for SC and 5.0 (± 0.6) for SF. Average difficulty rating per session were 6.4 (± 0.7) and 4.7 (± 0.9) for SC and SF respectively. A negative correlation (r=0.46, p*<* 0.01) was identified between the difficulty rating of the subjects and the block performance.

#### 3.2.4 Audio Stimuli

Finally we investigated the influence of the stimuli on the AAD performance measures in the current study. Due to the longitudinal setup of the study (i.e. *>*3 hours of unique audio stimuli, with 32 blocks per subject) it allows to look for effects that are related to the audio stories used. Both subjects were presented with the same order of audio stimuli. Posthoc we investigated the correlation between accuracy, CA-CUA mean and variance of the blocks, between both subjects. Figure 9 shows the block performances for both subjects throughout the experiment. A significant correlation was identified between the two subjects for block performances (r=0.46, p=0.009) and for mean CA-CUA (r=0.39, p=0.029, not shown in figure). In contrast, the CA-CUA variance did not correlate significantly between our subjects (r=0.24, p=0.179). Furthermore, we found a moderate correlation for mean CA (r=0.52, p=0.002), while mean CUA did not correlate significantly (r=0.25, p=0.164). These values indicate a dependency on the stimulus (i.e. story in each block) in which the AAD performance of both subjects were affected in similar ways.

**Figure 9:**
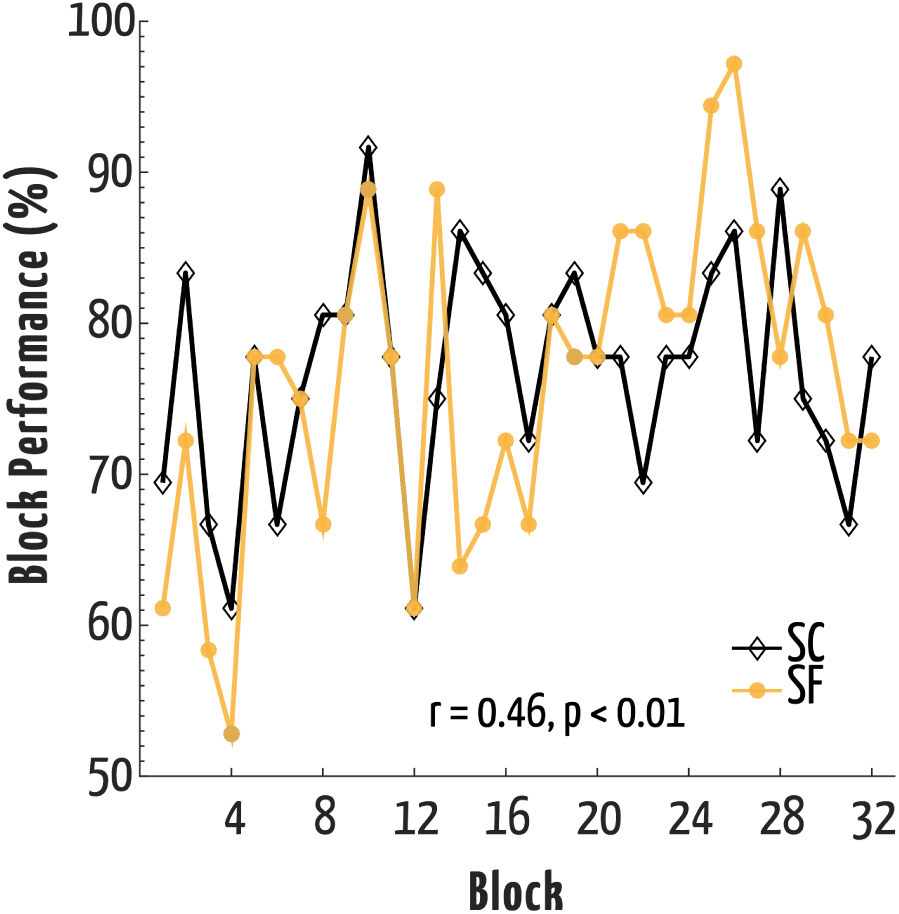
Block performances for both subjects and correlation between both subjects (r=0.46, p=0.009).

## 4 Discussion

In the current study, we evaluated the possibility of implementing an online closed-loop system for auditory attention detection and investigated the potential of providing feedback of the AAD performance to the subject. With window lengths of 10s, we obtained robust accuracies by detection auditory attention. We provided a fully working feedback system, and the online implementation did not significantly degrade the results. In the first study a slight improvement in accuracy was observed when using neurofeedback; although the number of subjects was limited it is the first study reporting such effects. In the second study a more realistic scenario was established for long-term training; we evaluated the effects on various parameters of the AAD during training with feedback for several days. Changes were noted for the subject that received feedback training which had a positive effect on its AAD accuracy. The effect was mostly caused by a more stable neural response of the experimental subject to the auditory attention task (i.e. a more consistent CA-CUA). Due to the nature of our study, it cannot be determined with full clarity to what extent these effects were caused by true self-regulation of neural activity of the experimental subject; or whether these effects were caused by an increased motivation or effort of the experimental subject compared to the control subject. Overall, the two studies we presented achieved results that were similar and competitive to existing lab-studies, especially since we used mobile EEG equipment and natural recordings settings (i.e. office and home). This is particularly interesting for future application in other (long term) studies in real-life conditions. Moreover, the detected changes in AAD for the feedback subject during and after training (study II) suggest that changes in AAD may be achieved via training. Future studies should unravel the causes of these changes in more detail to pave the way for improved AAD. Finally, we revealed a significant relationship between the audio stimuli and the performance of the two subjects. Future studies should (re)examine and choose their audio stimuli with care to avoid the heterogeneity between speech stimuli observed in our Study II. A quantifiable approach to handle the stimuli or shared audio database could help in comparing between AAD studies.

The presentation of neurofeedback cues on screen showed overall, in both studies, no negative deflection in accuracy due to increased cognitive demands being in a dual-task scenario. For example, the visual cues could lead to increased distraction. Nevertheless, for the first study, subjects who saw light-red feedback were more prone to perform bad in the next trial as well. One explanation may be that when subjects became aware of an error (i.e. shift from green trial to light-red) this leads to a brief surprise effect which lowers the attentional response in the next 10 seconds. On the contrary, after seeing the worst option (dark red) subjects were less likely to perfom bad on the next trial. This may have triggered an increased attentional effect or focus to perform better (since at the moment the subject cannot score worse). Faster feedback may be possible considering that at a trial length of 5 seconds above 70% accuracy can be obtained. This would mean that the subject sees twice as many feedback cues which may increase the potential of finding a true feedback training effect because the changes (and thus control of the training) are more instant. Alternatively, a sliding window approach (e.g. evaluate CA-CUA in the past 10s every *x* seconds) would allow more frequent updating of the feedback cue. In addition, a paradigm in which the subjects switched more frequently the attentional side (i.e. left and right) mimics more closely a real-world application albeit more difficult for the user. This increased difficulty may increase the reliance on the neurofeedback. This factor is not well reflected in the current paradigm; subjects only switched attention after each story.

The true mechanism behind the changes of the AAD performance of the subjects that received online feedback remains up to speculation. Besides the aforementioned results and explanation, there is another area of change that may have occurred but remains hidden in the experiment; users may have adapted (un)consciously their listening behavior. In other words, different listening strategies, for example repeating words silently while listening, or focus particular on silences in between words. Experimental results of such changes are non existing to date and would be very interesting to investigate as such changes would be highly valuable for future users of auditory-assistive devices as these behavioral attention effects might boost the AAD analysis accuracy. Interestingly, the difficulty rating in both studies correlated to performance on subject (Study I) and block level (Study II). This might indicate that specific aspects of the stories lead to in- or decreased detection accuracy which would be an interesting line of future research.

The high accuracies in the present work were obtained with 24 electrodes and were found to be stable (not significantly different) up to removal of 16 channels (not presented in the results section). These results are in line with insights presented by [26] and are encouraging for future work in online processing and mobile EEG experiments. It would be particularly interesting to see an online implementation as presented here to be combined with the around- or in-the-ear electrodes as recently described in [25]. Moreover, meaningful accuracies were obtained even at window lengths as low as 3 seconds which opens up possibilities for fast feedback. To this end, it would also be interesting to see how these results relate to a state-space model, as the latter was shown to have a high temporal resolution in MEG recordings [2]. It is interesting that an increased AAD trend remains after training in Study II. Up to 10 days later, this trend could be noted in accuracy and variance of the signals (note that these effects seem to decrease from FU1 to FU2). The evaluation of neurofeedback effects lack follow-up sessions after a few weeks or even months. In real-life we would want long lasting (permanent) changes of the subject in relation to the AAD. In many neurofeedback studies the large amounts of recording sessions is seen as a hurdle. Nevertheless the option to use mobile EEG hardware facilitates more easily and convenient future recordings to overcome these limitations.

A general effect of performing repetitive AAD blocks, regardless of the presentation of feedback, could not be established due to the influence of the story stimuli. Being the first longitudinal AAD study many more stimuli were used than previously reported in literature. This is likely to have caused increased variation in our stories that in turn influenced the performance. The significant correlation between the two subjects indicates these changes (i.e. in story content and/or acoustic features). They are not caused by a general increase or decrease (trend) as is evident from the highly scattered pattern in time in Figure 9A. Such variations in stimuli are not present in other auditory paradigms such as the P300 oddball (i.e. pure tones), or syllables without semantic meaning [18]. The latter holds potential to obtain complementary information on the segregating process of speech mixtures into distinct streams with respect to AAD studies that rely solely on natural speech stimuli. For AAD our results on the audio stimuli raise the question to what extent the stimuli in existing publications in the domain can be related to each other and maybe account for differences between studies (or within). Therefore we suggest to look into ways of quantifying more precisely the stimuli used for AAD. One influential factor may be the window length in RMS normalization. Shorter windows may lead to smaller differences between the two speech waveforms although it remains up to speculation to what these differences can be attributed; future work could look into the complexity and content of the stories, emotion levels, personal preference of the subject, among others. Another solution may be the sharing of stimuli, although most studies use stimuli in their subjects’ native language, which makes this more difficult.

Another limitation in the current methodology is the lack of incorporating real-life audio signals. Recently there have been studies evaluating the effect of noisy reference signals and showing a negative impact on performance [35, 6] Another aspect that remains unclear is whether the spatial location of the speaker will influence the classification accuracy and, to what extent a feedback procedure is reflected in the detection with multiple speakers. Finally, the nature of study II, being a case study with only two subjects, limits the generality of the results to certain extent. Nevertheless, promising results were obtained even in the limited setup paving the way for new experimental studies in the field of AAD.

All in all, we showed (to the best of our knowledge) the first closed-loop AAD system with a rather convenient recording setup and hardware. The described results are very promising to conduct future online experiments with (mobile) EEG hardware in order to obtain more robust conclusions about increasing AAD performance of the subjects through online feedback. Especially to investigate the effects of real-life distractions (e.g. cognitive load [41]) on the AAD in a cocktail party scenario would be very valuable as the brain signals in such scenarios may differ diminishing the application potential of existing AAD systems.

## 5 Conclusion

In the current study, we have implemented a fully automated closed-loop system that allows for convenient recording outside a lab environment to detect auditory attention in a dual speaker scenario. We achieved high auditory attention detection accuracies with a trial length of 10 seconds and provided subjects with online visual feedback on their ongoing performance. This exploratory study proves the feasibility of investigating the effect of neurofeedback in such a setting, paving the way for future studies.

Observed changes in the AAD have been linked to the presentation of the feedback process. Moreover these effects were observed a couple of days later as well. The degree of which this is related to the neurofeedback, increased motivation or attentional effort remains to be unraveled. Nevertheless, these results suggest that the AAD in healthy subjects is versatile (to changes). Future work should therefore focus on addressing the cause of these changes in more detail and opt for universal audio (speech) stimuli. If these changes can be made, AAD training could be highly beneficiary for future users and increase the application potential for subjects that seem to be less able to use the AAD paradigm at the moment.

## Acknowledgements

A conference precursor of this manuscript has been published in [40]. Research supported by Research Council KUL: CoE PFV/10/002 (OPTEC); Belgian Federal Science Policy Office: IUAP P7/19/(DYSCO, 20122017), Flemish Government, FWO projects: G.0427.10N; EU: European Research Council under the European Union’s Seventh Framework Programme (FP7/20072013)/ERC Advanced Grant: BIOTENSORS (nr. 339804). This paper reflects only the authors’ views, and the Union is not liable for any use that may be made of the contained information.

